# SC-MAMBA2: Leveraging State-Space Models for Efficient Single-Cell Ultra-Long Transcriptome Modeling

**DOI:** 10.1101/2024.09.30.615775

**Authors:** Yalong Zhao, Bowen Zhao, Fan Zhang, Chenfeng He, Wendao Wu, Lipeng Lai

## Abstract

The rapid advancement of single-cell sequencing technology has significantly deepened our understanding of cellular heterogeneity, yet it concurrently presents substantial challenges for the unified modeling of single-cell data. Simultaneously, pre-trained foundation models have achieved notable success in domains such as natural language processing and image analysis. However, extending these models to accommodate ultra-long single-cell transcriptome sequences, characterized by an extensive number of genes, remains a formidable task. In this study, we introduce SC-MAMBA2, based on the MAMBA2 architecture, meticulously designed with a bidirectional modeling approach tailored for single-cell transcriptomics data. As the first single-cell foundation model to integrate state-space models (SSMs) underlying MAMBA2 architecture, SC-MAMBA2 features over 625 million parameters, covers more than 60,000 genes, and was pre-trained on a dataset of over 57 million cells, making it the most comprehensive solution for processing ultra-long transcriptome sequences. Extensive bench-marking across a diverse array of downstream tasks consistently demonstrates that SC-MAMBA2 surpasses state-of-the-art models, delivering superior accuracy and enhanced computational efficiency.

## 1 Introduction

The paradigm of large-scale pretraining followed by fine-tuning on specific datasets has significantly advanced computational biology, making the use of foundation models for biological data analysis a growing research trend [1, 2, 3, 4]. In the single-cell field, various foundation models have already been developed[5, 6, 7, 8, 9]. By pretraining on extensive gene expression datasets, these models gain an understanding of complex, high-dimensional gene regulatory relationships and perform well on downstream tasks after fine-tuning, all while avoiding the need for cumbersome feature engineering and reducing manual intervention during data analysis. Overall, single-cell foundation models have facilitated the elucidation of cellular heterogeneity and the comprehension of complex biological systems[5], driving the progress of precision medicine in the single-cell domain.

Given the complex regulatory mechanisms between single cells, the attention mechanism[10] has become an effective way to capture relationships between genes. Most foundational single-cell models are also built upon attention mechanisms. scBERT [8], based on the BERT[11] model, employs linear attention[12] to perform masked modeling across all genes, enhancing the performance of single-cell annotation. GeneFormer converts the absolute expression levels of each gene into relative rankings and uses masked learning to understand gene interactions and regulatory networks. scFoundation infers masked genes through unmasked genes, introducing the masked auto-encoder[13] framework into single-cell modeling, and demonstrating its ability to enhance single-cell representations as a standalone module. scGPT first learns via self-attention, then applies cross-attention[10] to predict unknown genes, which improves the performance of various downstream tasks.

However, the aforementioned models still face some inherent limitations. Most foundational single-cell models rely on masked learning and have yet to tap into the vast potential of generative models[1, 2, 3, 4]. Additionally, the quadratic complexity of attention calculations leads to significant resource consumption during training and inference, making it challenging to scale to larger datasets, as seen in fields like natural language processing[14, 15]. As a result, exploring how to introduce efficient language models capable of modeling complex gene dependencies into single-cell data analysis has become a promising new opportunity. Mamba [16], inspired by classical state space models, offers near-linear scalability with respect to sequence length while retaining modeling capabilities comparable to Transformers. This makes it an emerging framework in the natural language processing domain[17], and it has already seen active applications across various fields, including computer vision[18] [19], natural language processing [20] [21], and healthcare [22] [23] [24].

In this work, we introduce SC-MAMBA2, a generative foundational model for single-cell data analysis with over 625 million parameters, pre-trained on a dataset containing 57 million cells. Our backbone network leverages Mamba2[25][16] to achieve fast inference and linear scaling of feature dimensions, efficiently modeling interactions between tens of thousands of genes while simultaneously capturing representations of both cells and genes. Additionally, we designed a bidirectional modeling approach tailored to the non-sequential nature of single-cell data, making SC-MAMBA2 a generative pre-training foundational model specifically for single-cell transcriptomics data. We also adapted SC-MAMBA2 to various downstream tasks, enhancing the utility of the pre-trained model across a wide range of applications.

The contributions of this work are threefold:

### Innovative Architecture

Innovative Architecture: We introduce SC-MAMBA2, the first model to integrate state-space models (SSMs) with the MAMBA framework for single-cell ultra-long transcriptome data. This integration enables efficient and scalable modeling of large gene sequences, overcoming the computational efficiency limitations of traditional Transformer-based architectures when handling large-scale biological data.

### Design Adaptations and Long Sequence Modeling

We develop unique design modifications tailored to gene sequences and implement a bidirectional modeling approach within the state-space modules. SC-MAMBA2 is capable of modeling full gene sequences encompassing 60,530 genes, representing the largest and most comprehensive sequence length handled in the single-cell transcriptomics domain to date. This capability allows for comprehensive analysis of entire gene transcripts, capturing intricate biological variations and regulatory elements that shorter models cannot accommodate. By successfully modeling such extensive sequences, SC-MAMBA2 provides a more complete and accurate representation of the transcriptome, leading to superior performance in downstream tasks and setting a new benchmark for future studies in the field.

### Robust performance

Through extensive benchmarking, we demonstrate that SC-MAMBA2 outperforms existing state-of-the-art models across multiple downstream tasks, including gene expression quantification, cell type classification, and trajectory inference. This superior performance underscores SC-MAMBA2’s efficacy in capturing the full breadth of transcriptomics information while maintaining computational efficiency, contributing to the broader adoption of foundational generative models in transcriptomics.

By addressing the computational challenges inherent in single-cell ultra-long transcriptome modeling, SC-MAMBA2 paves the way for more comprehensive and efficient analyses, ultimately contributing to a deeper understanding of the mechanisms driving complex biological systems and advancing single-cell data analysis towards foundational models.

## 2 Results

SC-MAMBA2 has over 625 million learnable parameters and has been pre-trained on over 57 million cells from the CELLxGENE [26] dataset, covering more than 60,000 genes. It is based on MAMBA2 and has been redesigned with a bidirectional architecture to adapt to the unique characteristics of single-cell data, enabling efficient learning of dependencies between all genes. The model offers extensive gene context length, coverage, and scalability, enhancing the generative foundation for single-cell data analysis. After fine-tuning for specific tasks, SC-MAMBA2 has outperformed benchmark methods in cell annotation, multi-batch data integration, multi-omic data integration, and perturbation prediction. The results demonstrate that the model accurately captures complex gene dependencies and provides a robust pipeline for downstream analysis.

### 2.1 SC-MAMBA2 improves multi-batch and multi-omic integration

#### Multi-batch integration

Due to the existence of multiple platforms and diverse sequencing protocols[27, 28], batch effects between single-cell sequencing data are inevitable. Therefore, to gain unified insights from single-cell data of the same tissue, it is necessary to integrate and analyze data from different batches. This also poses a challenge for batch integration algorithms in eliminating batch effects while preserving true biological variations. We used the [CLS] token to represent each cell, followed by an adversarial domain adaptation approach to remove batch effects between cells. In our benchmarking tests, we compared SC-MAMBA2, scGPT, and three popular integration methods: scVI[29], Seurat[30], and Harmony[31]. In the PBMC 10K dataset[32], SC-MAMBA2 successfully distinguished all cell types, achieving an AvgBIO score (the mean of NMI, ARI, and ASWcell[33]) of 0.836. This represented a 7%-12% improvement over scVI, Seurat, and Harmony, and a 1.83% improvement over scGPT, demonstrating the robust batch integration capability of SC-MAMBA2(Fig 2a, Fig2d).

**Figure 1:**
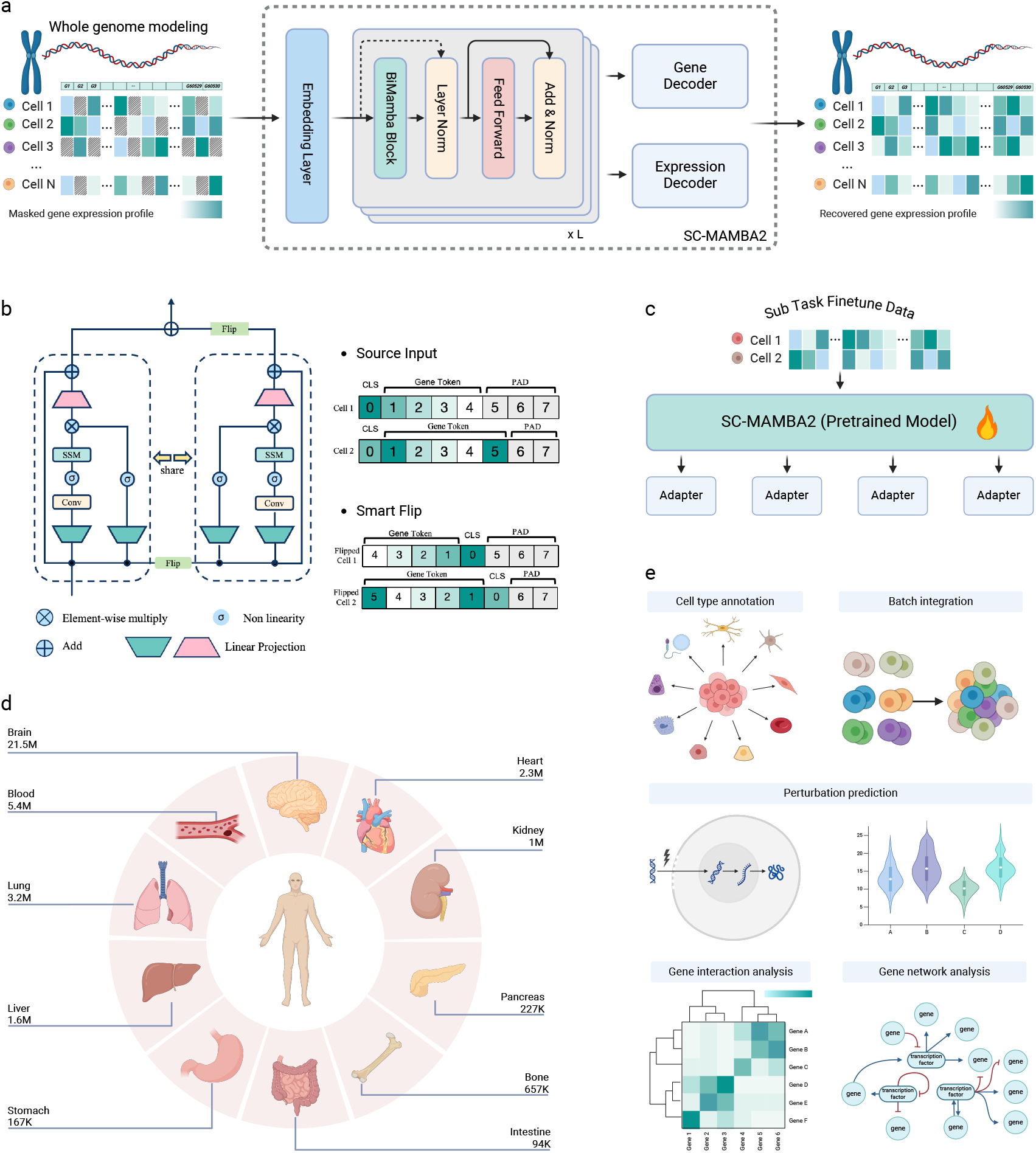
Schematic overview of the SC-MAMBA2 architecture and its applications in single-cell transcriptomics. **a**, Whole genome modeling pipeline using SC-MAMBA2. The masked gene expression profiles from multiple cells are processed through the SC-MAMBA2 architecture. The output is then passed to both Gene and Expression Decoders, recovering the gene expression profile of each cell. **b**, Smart Flip bidirectional input mechanism using state-space models (SSMs). The architecture supports both source input, where the gene tokens are processed in a specific order, and a Smart Flip mechanism, where the sequence is reversed, enhancing bidirectional transcriptome modeling. **c**, Adapter-based fine-tuning of the SC-MAMBA2 pre-trained model for sub-task adaptation using specific data. **d**, Extensive and diverse training data sourced from CELLxGENE, covering multiple organs and tissues. **e**, Downstream applications of SC-MAMBA2, including cell type annotation, batch integration, perturbation prediction, gene interaction analysis, and gene regulatory network inference, showcasing its versatility in various single-cell tasks.

**Figure 2:**
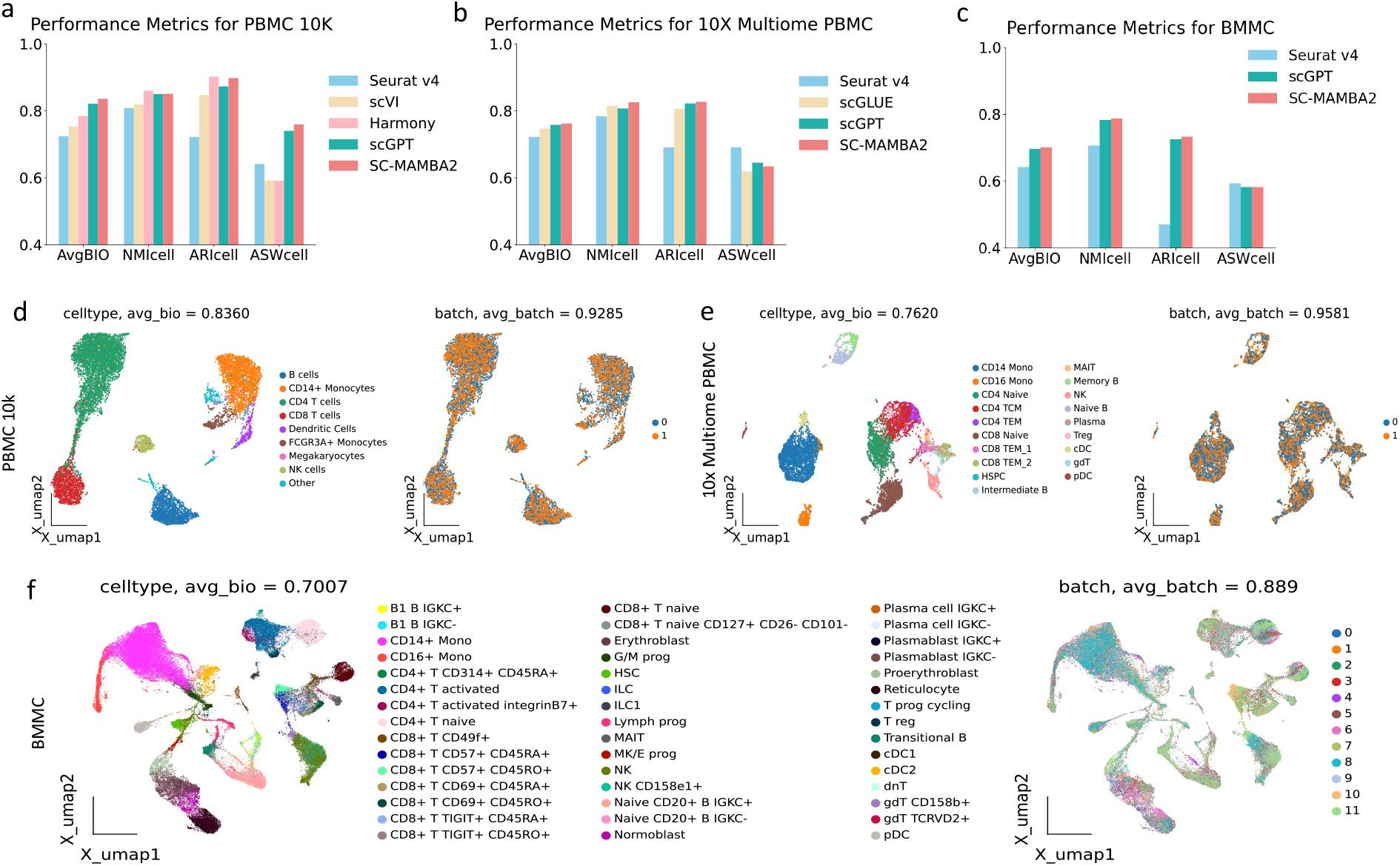
SC-MAMBA2 performance on multi-batch and multi-omic integration. **a**, Comparison of batch integration performance among Seurat, scVI, Harmony, scGPT, and SC-MAMBA2 on the PBMC 10K dataset. **b**, Comparison of integration performance among Seurat, scGLUE, scGPT, and SC-MAMBA2 on the 10X multi-omics PBMC dataset. **c**, Comparison of integration performance among Seurat, scGLUE, scGPT, and SC-MAMBA2 on the BMMC dataset **d**, The UMAP of fine-tuned SC-MAMBA2 on the PBMC 10k dataset for the cell type-clustering and batch integration task. The UMAP plot of learned cell embeddings is colored by cell types (left) and different batches (right). **e, f**, The UMAP visualization of fine-tuned SC-MAMBA2 on the PBMC 10K multi-omic dataset (e) and BMMC dataset (f), showing cell types (left) and cell modalities (the BMMC dataset includes two modalities and multiple batches) (right).

#### Multi-omic integration

Single-cell multi-omics provides multiple perspectives on observing cell states. For example, scATAC-seq focuses on chromatin accessibility, scRNA-seq monitors transcription levels, while single-cell proteomics examines the abundance of translated proteins. Consequently, understanding various aspects of cell states using multiomic data has become a current research hotspot. The 10X Multiome PBMC dataset[34], which includes measurements of gene expression and chromatin accessibility, was used to benchmark SC-MAMBA2, scGPT, scGLUE[35], and Seurat (v.4)[30]. SC-MAMBA2 demonstrated more clearly defined clusters compared to scGPT. Although the improvement in AvgBIO score was only 0.5%, SC-MAMBA2 was the only method to successfully differentiate CD14+ and CD16+ naive cells into distinct clusters (Fig 2c, Fig 2e). In the bone marrow mononuclear cells (BMMCs)[36] dataset, which contains gene expression and protein abundance data, including 90,000 cells, 12 batches, and 48 cell types, SC-MAMBA2 showed significant performance improvements compared to scGPT and Seurat (Fig 2c, Fig 2f).

### 2.2 SC-MAMBA2 performance on cell type annotation

There are now several comprehensive cell atlases for various tissues and organs[37, 38], and transferring the detailed cell annotation labels from these annotated single-cell datasets to unannotated datasets will greatly simplify the single-cell analysis workflow, enabling accurate annotation of hard-to-distinguish cell subtypes. We conducted supervised fine-tuning of the model using a reference cell atlas containing ground truth labels and evaluated the annotation performance using an independent query dataset. SC-MAMBA2 does not filter gene expression data but retains all genes, performing normalization, log transformation, and binning. The model then infers the [CLS] token as a representation of each cell. Additionally, we incorporated an extra classifier to predict cell type labels and minimized classification loss during training.

We conducted supervised fine-tuning of the pretrained SC-MAMBA2 and applied it to three datasets: hPancreas[39], Myeloid[40], and Multiple Sclerosis[41], benchmarking it against two other Transformer-based methods, TOSICA[39] and scGPT. The results indicate that SC-MAMBA2 achieved equal or superior accuracy, precision, recall, and macro F1 scores[42] across all cell annotation tasks compared to the other models (Fig. 3d). In both the hPancreas and Myeloid datasets, SC-MAMBA2 demonstrated improvements in precision, recall, and macro F1 scores compared to scGPT, despite similar accuracy levels. Figures 3a, 3b, and 3c present the predicted cell annotations using SC-MAMBA2 after fine-tuning, showing its ability to accurately identify most cell types. This suggests that the cell representations extracted by SC-MAMBA2 effectively capture cell type-specific signals.

**Figure 3:**
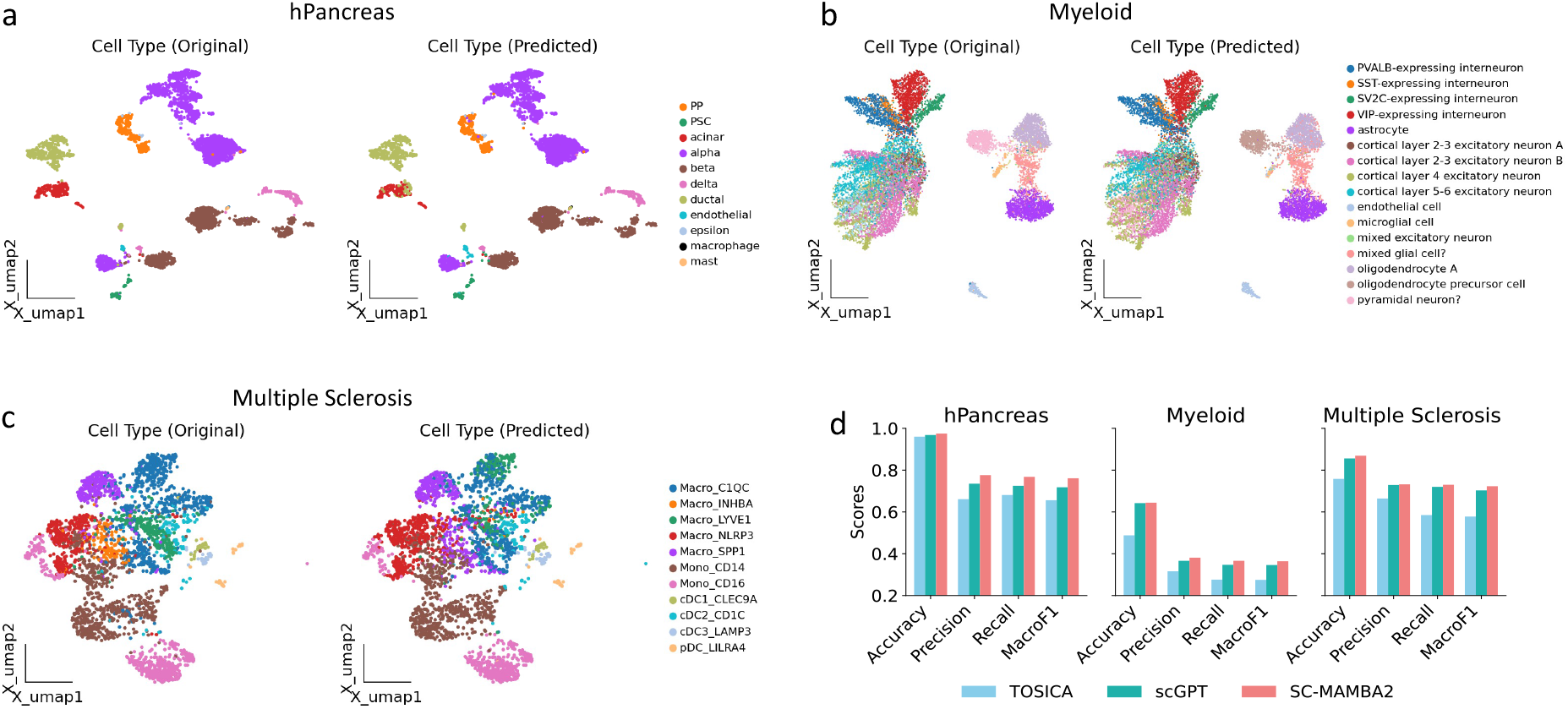
Cell type annotation with SC-MAMBA2. The UMAP displays the cell annotation results for the human pancreas dataset (**a**), myeloid cells (**b**), and multiple sclerosis (**c**), colored by the cell types annotated in the original study (left) and by the cell types predicted by the fine-tuned SC-MAMBA2. (right). **d**, The barplot of accuracy, precision, recall and macro f1 score compared between SC-MAMBA2 and TOSICA, scGPT in three datasets.

### 2.3 SC-MAMBA2 improves In silico perturbation and reverse perturbation prediction

Gene regulatory networks are hierarchical, where a change in the expression of one gene can influence the expression of multiple other genes [43]. Predicting how other genes’ expression levels respond to perturbation of a single gene can help uncover the hierarchical structure of these networks. Conversely, identifying key genes that drive changes in gene expression under specific conditions, such as diseases (i.e., reverse perturbation), can inform the development of therapies targeting the core regulatory elements responsible for the disease, rather than focusing on downstream effects that may not be directly related to the disease [6].

In this study, we tested SC-MAMBA2 and benchmarking methods on perturbation prediction using the Norman dataset [44], which includes 105 single-gene perturbations and 131 two-gene perturbations. Specifically, we utilized a subset of perturbation data (Fig. 4c), adding perturbation tokens to the original embeddings and inputting them into the pre-trained models. The models were fine-tuned in a supervised manner to predict post-perturbation gene expression, aligning the predictions with the actual measured data. Given the involvement of two-gene perturbations, we evaluated the models under four experimental conditions: “single unseen” for predicting unseen genes in single-gene perturbations, and “seen 0,” “seen 1,” and “seen 2” representing the number of observed genes in two-gene perturbations. SC-MAMBA2 consistently outperformed other methods, achieving at least a 5% higher Pearson correlation with measured gene expression changes (Fig. 4a). Notably, SC-MAMBA2 excelled in two-gene perturbations, especially when neither gene had been seen during fine-tuning (Fig. 4b), indicating its superior generalization ability for novel perturbed genes.

**Figure 4:**
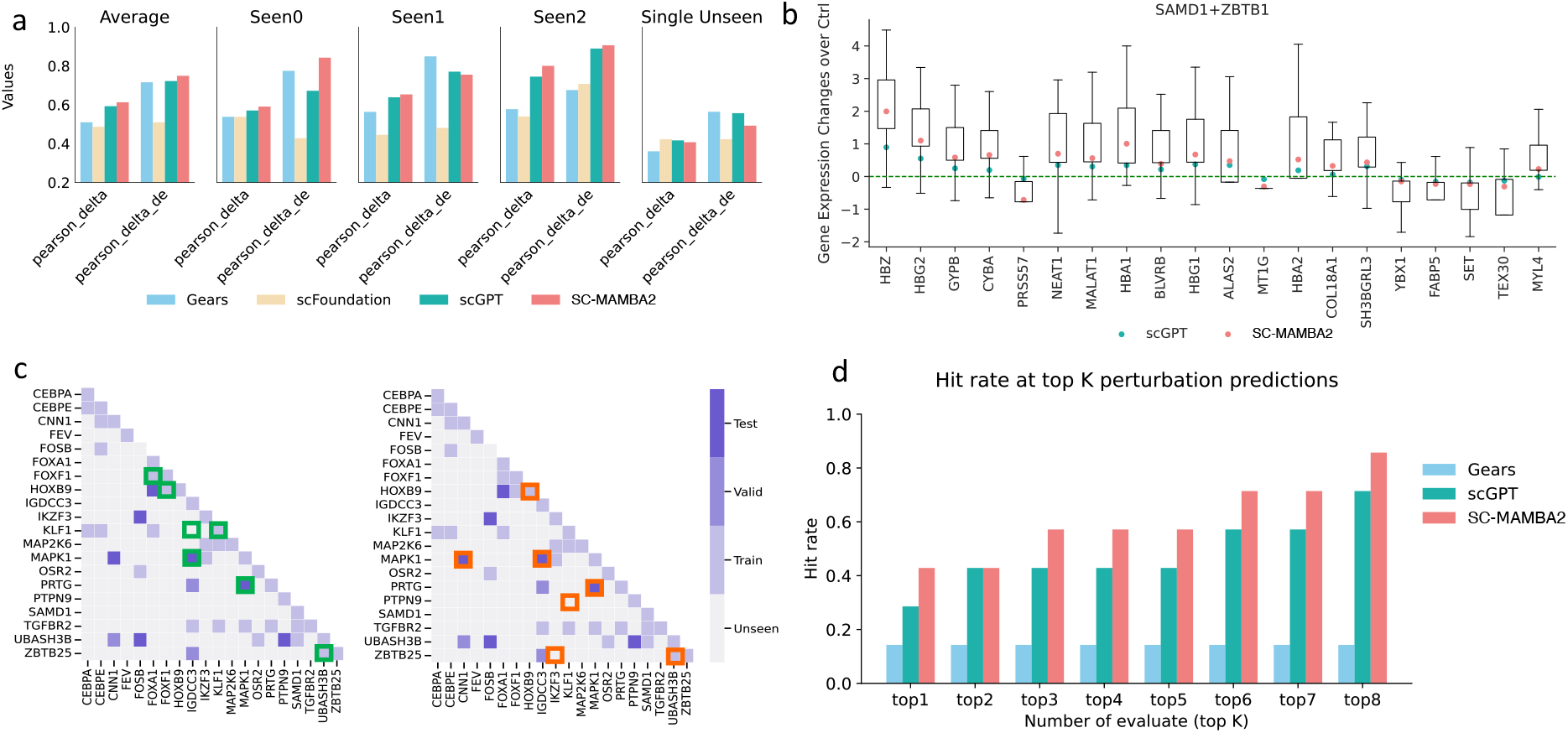
Evaluation of models accuracy for perturbation and reverse perturbation prediction. **a**, SC-MAMBA2 scGPT and GEARS[45] performance on perturbation task. Perturbations in the test dataset were separated into 4 categories: Single Unseen (single-gene perturbed and this gene was not in the fine-tune training dataset), Seen0 (two-genes perturbed and both genes were not in the fine-tune training dataset), Seen1 (two-genes perturbed and one of the genes was in the fine-tune training dataset) and Seen2 (two-genes perturbed and both genes were in the fine-tune training dataset). The averaged metrics were also shown (left). Two metrics were included for evaluating the accuracy: pearson_delta (pearson correlation between the predicted versus observed gene expression after subtracting the control cells’ gene expression profile) and pearson_delta_de (pearson_delta of the all the differentially expressed genes) **b**, Example perturbation (SAMD1+ZBTB1) prediction results, showing the distribution of predicted and observed gene expression for the top 20 differentially expressed genes. The box denotes the interquartile range of expression change. The green dot represent the prediction value from scGPT and the red dot represent the value predicted by SC-MAMBA2. **c**, Visualization of scGPT and SC-MAMBA2 reverse perturbation prediction results. Heatmap show the possible perturbation combinations. Grid color represents the experiment type (train, valid, test and unseen). Green square boxes are scGPT prediction results (left) and red boxes are SC-MAMBA2 results (right). **d**, Comparison of Reverse Perturbation Accuracy Between scGPT and SC-MAMBA2. The hit rate was defined as the number of correct perturbations accurately predicted within the top K (1 to 8) rankings.

Predicting the cause of perturbation from the perturbed state also helps us understand disease progression and identify potentially meaningful therapeutic targets. Using the same reverse perturbation setup as in scGPT[6], we aimed to trace the perturbed genes from the perturbation states. The experimental results show that SC-MAMBA2 outperformed scGPT in terms of accuracy. By exploring the combination space of the top 20 expressed genes (Fig. 4c), SC-MAMBA2 accurately identified three out of seven unseen perturbations in its top 1 prediction, whereas scGPT only correctly predicted two. When extending the evaluation to the top 1-8 predictions, SC-MAMBA2 consistently outperformed both scGPT and GEARS in the number of accurate predictions (Fig. 4d).

### 2.4 SC-MAMBA2 uncovers gene network of specific biological functions

Genes function collaboratively, working as networks to facilitate essential biological processes. scGPT showed its ability for gene network analysis (Fig. 5a). Here, we demonstrate the pretrained SC-MAMBA2 model embedding can also capture the biological relevance among genes. To do this, we extracted the embedding of all the CD genes (n=80) from SC-MAMBA2 pretrained model (without fine-tuning, Fig. 5b). Cosine similarity was employed to assess the similarity of the embedding vectors. Subsequently, Leiden clustering was used to group genes with similar embeddings into four distinct clusters (Fig. 5c). Interestingly, each cluster was dominated by genes of different functions (Fig. 5d). Specifically, all clusters were notably enriched with CD genes associated with the immune system, which aligns with expectations. Besides, genes in Cluster 3 (orange) are predominantly related to innate immune functions, while Cluster 1 (blue) is more enriched with genes involved in adaptive immune functions. Cluster 0 (red) comprises a mix of genes related to both innate and adaptive immunity, spanning across the gene network and pathways. Cluster 2 (green) is primarily enriched with genes related to B-cell receptor (BCR) functions. In summary, the gene embeddings from the pretrained SC-MAMBA2 model effectively capture the specific biological functions of different genes.

**Figure 5:**
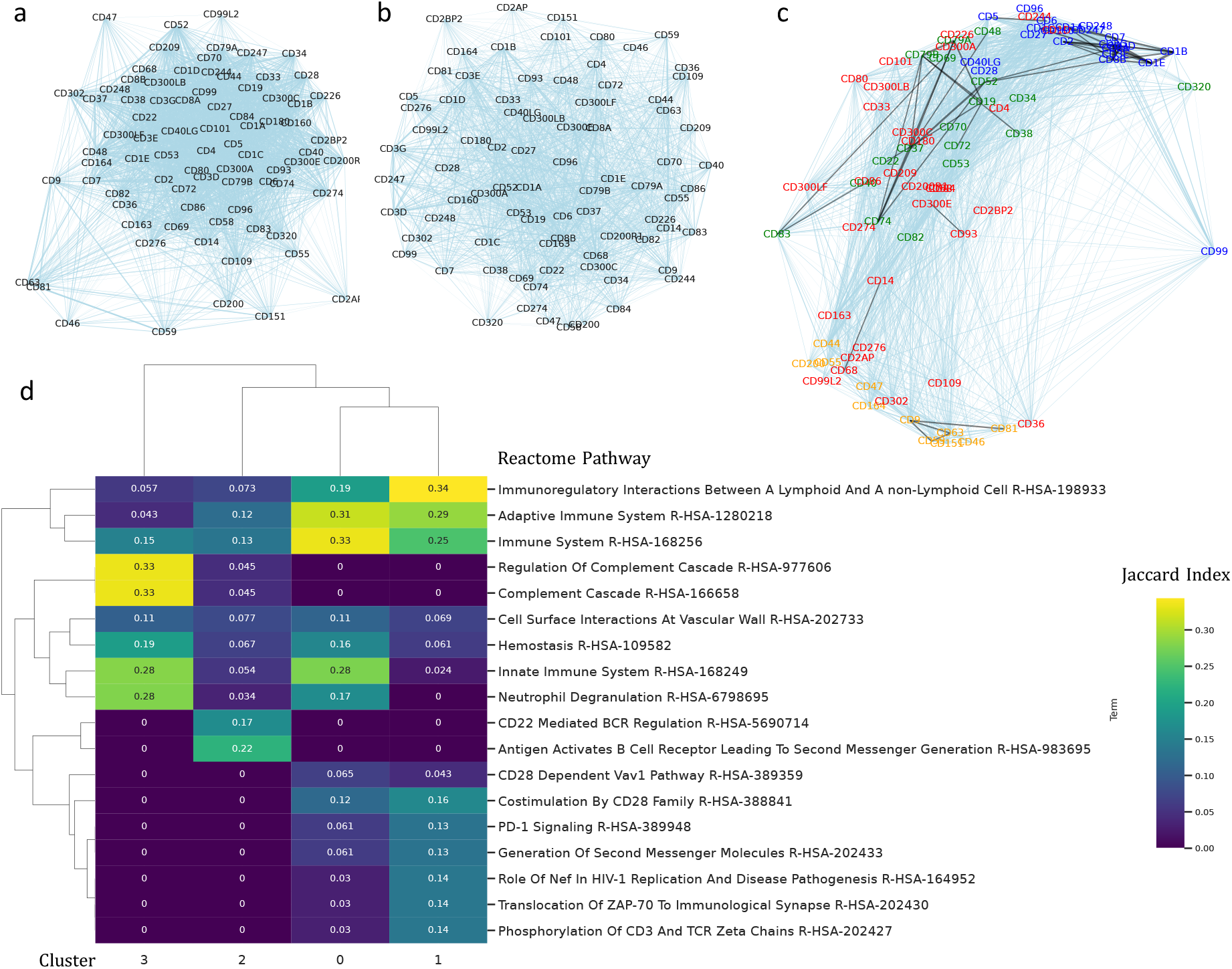
SC-MAMBA2 model for gene network analysis. **a**, Gene network predicted from scGPT embedding. Edges weight were measured as the cosine similarity of the genes embedding. **b**, Gene network predicted from SC-MAMBA2 embedding. **c**, Genes within the SC-MAMBA2 network were grouped into 4 clusters (cluster0: red, cluster1: blue, cluster2: green, cluster3: orange) based the network edges and connectivity. **d**, Functional analysis of each cluster by comparing the Jaccard Index of overlapping genes between each cluster and each Reactome pathway (https://reactome.org/[46]).

## 3 Discussion

In this study, we developed SC-MAMBA2, the largest foundation model to date for modeling single-cell transcriptomics. The efficiency of the Mamba2 algorithm, combined with our unique model design for transcriptomics data, enables SC-MAMBA2 to more effectively and accurately model complex single-cell gene regulatory networks, learning superior cellular representations. Through tasks such as batch integration, multi-omic integration, cell annotation, perturbation modeling, and gene interaction analysis, we demonstrated that large-scale generative foundation models excel in various single-cell downstream tasks, outperforming several state-of-the-art single-cell foundation models. This accelerates the efficient analysis of single-cell data and provides potential therapeutic targets for real diseases. In the future, we plan to incorporate more gene perturbation and drug perturbation data to construct disease models, and combine organoid drug screening to advance personalized treatment strategies for different disease subtypes.

## 4 Methods

### 4.1 State Space Models

State Space Models (SSMs) are traditionally used to describe the dynamics of continuous systems by transforming an input sequence *x*(*t*) ∈ ℝ into a latent state representation *h*(*t*) ∈ ℝ^*N*^. This latent state is then utilized to generate an output sequence *y*(*t*) ∈ ℝ. The mathematical formulation of an SSM is given by:

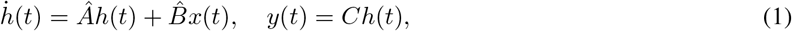

where *Â* ∈ ℝ ^*N×N*^, 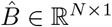, and *C* ∈ ℝ ^1*×N*^ are parameters. To effectively incorporate continuous SSMs into deep learning architectures, discretization is essential. This involves introducing a time step parameter Δ ∈ ℝ and applying the zero-order hold (ZOH) technique for discretization.

Through this process, the continuous matrices *Â* and 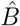 are transformed into their discrete counterparts, *A* and *B*. Consequently, Equation (1) is reformulated in discrete form as shown in Equation (2), making it more suitable for implementation in modern computational frameworks:

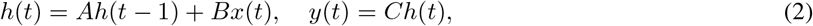

where 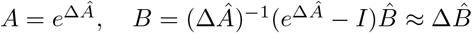, and *I* denotes the identity matrix. Additionally, the process described in Equation (2) can be implemented globally in a convolutional manner as:

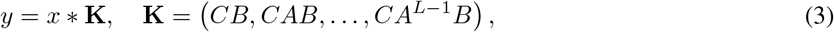

where **K** ∈ ℝ^*L*^ represents the convolution kernel. Recent work by Mamba proposes a method in which the parameters *B, C*, and Δ are input-independent, addressing limitations inherent in previous Linear Time Invariant (LTI) SSM models. This enhancement improves the adaptability and performance of SSMs.

#### State Space Duality

Mamba2 recently introduced the concept of State Space Duality (SSD), simplifying the matrix **A** into a scalar. This specific case of selective State Space Models (SSMs) can be applied in both linear and quadratic forms. Without loss of generality, the matrix transformation form of selective state space models is represented as follows:

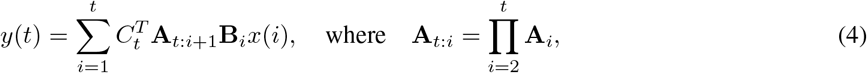

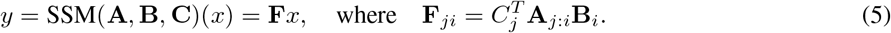

When *A*_*i*_ is reduced to a scalar, the quadratic form of Equation (4) can be reformulated as:

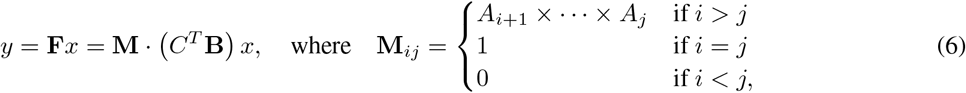

while its linear form is expressed as:

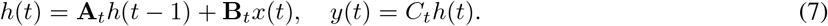

### 4.2 SC-MAMBA2

#### Pre-processing

In the data preprocessing stage, we first perform expression level statistics on the data. The number of genes expressed by each cell is denoted as *N*_*i*_. To ensure uniform input dimensions for batch processing in neural networks, we specify a maximum sequence length *M*. Depending on *N*_*i*_, we process each cell’s expression data as follows:

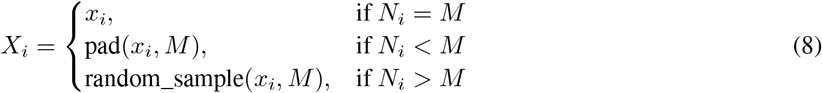

where *X*_*i*_ is the processed expression vector of length *M* for cell *i*, and *x*_*i*_ is the original expression vector.

The embedding of SC-MAMBA2 consists of two main parts: gene name encoding and expression value encoding.

#### Gene Name Encoding

Each gene *g*_*j*_ is tokenized and represented as a high-dimensional vector, similar to word embeddings in NLP. Specifically, each gene is indexed based on the dataset’s gene list (e.g., `CELLxGENE`) and encoded via a gene name encoder:

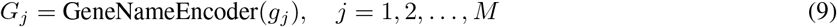

where *G*_*j*_ ∈ ℝ^*d*^ denotes the gene embedding in *d*-dimensional space.

This tokenization creates a dictionary that maps each gene name *g*_*j*_ to its corresponding vector, forming the set:

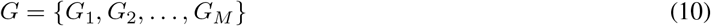

These embeddings are then utilized for downstream analysis, enabling a structured and learnable representation of gene names.

#### Expression Value Encoding

The expression values are binned according to their relative expression levels across cells. This binning process helps mitigate batch effects. Let *B* denote the number of bins. Each binned value *e*_*j*_ is then passed through an expression value encoder, which uses a continuous mapping to project the binned expression values into a *d*-dimensional space:

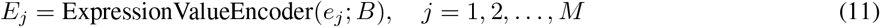

where *E*_*j*_ ∈ ℝ^*d*^ represents the encoded expression value in the *d*-dimensional space, and *B* indicates the total number of bins used in the binning process.

#### Cell Embedding

The embedding *C*_*j*_ for each gene in a cell is computed by adding the gene name encoding and the expression value encoding:

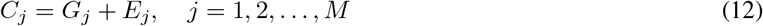

The complete embedding sequence for cell *i* is:

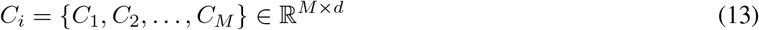

To facilitate the self-supervised training objective, we mask a certain proportion of the encoded expression values *E*_*j*_:

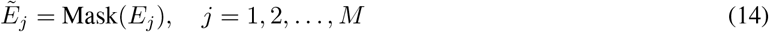

The model is then trained to predict the masked expression values 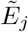 based on the context provided by the unmasked tokens. This masking mechanism allows the model to learn representations that generalize across different cells and genes.

#### Training

Since the original Mamba model is autoregressive and cannot capture bidirectional contextual relationships, we specifically designed the BiMamba module to process the embedding sequence *C*_*i*_. The input sequence *C*_*i*_ is first reversed to create a “flipped” sequence:

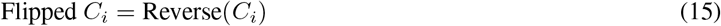

Both *C*_*i*_ and Flipped *C*_*i*_ are independently processed by weight-shared unidirectional Mamba modules:

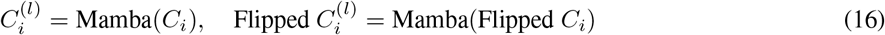

The flipped sequence is then reversed back to its original order:

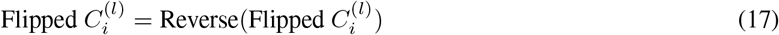

The outputs from both directions are combined through summation:

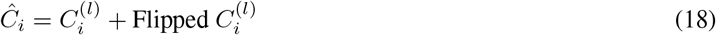

This combined sequence *Ĉ*_*i*_ is passed to the next layer in the BiMamba block. The Smart Padding mechanism ensures that only meaningful tokens (e.g., CLS token and gene tokens) are flipped, while padding tokens remain in their original positions to prevent artifacts during the reversing process.

Inspired by the Transformer architecture, conventional attention modules are replaced with BiMamba modules to construct a deep architecture capable of capturing bidirectional relationships. The final embedding sequence *C*_*i*_ is passed through a stack of *L* BiMamba layers, where the output of the *l*-th layer is given by:

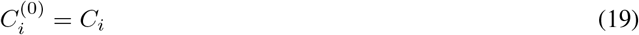

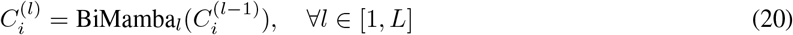

where 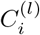 denotes the output of the *l*-th BiMamba layer. Each BiMamba_*l*_ builds upon the representation learned from the previous layer, and the output of each layer is used as the input to the next layer in the stack. This formulation ensures that the model progressively refines its representation, effectively capturing complex bidirectional relationships within the gene expression data.

#### loss

The gene expression prediction module processes each gene token to output the predicted binned expression value 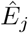. The loss function is computed as the mean squared error (MSE) between the predicted binned expression values 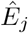 and the original binned expressions *E*_*j*_:

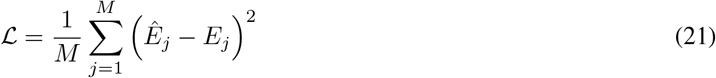

This loss function enables end-to-end training of the model, allowing it to learn to predict masked expression values based on the context. By stacking multiple BiMamba layers, the model is trained to capture complex bidirectional relationships within the gene expression data.

### 4.3 Cell type annotation

To address the cell type annotation task, we fine-tuned the pretrained SC-MAMBA2 using a reference dataset that included true labels and subsequently evaluated its annotation performance on an independent query dataset. We retained the gene markers shared between the pre-trained base model and the reference dataset. During fine-tuning, all gene markers were considered, regardless of whether their expression values were zero. Prior to fine-tuning the model, the gene expression values underwent normalization, log-transformation using Scanpy[47], and binning consistent with the pre-training process. The fine-tuned model was initialized with the weights of the pre-trained model, except for the cell type classifier, which was randomly initialized. During fine-tuning, the objective was to minimize the classification loss using a cell type classification fine-tuning approach. We used sklearn[42] to compute the supervised evaluation metrics. All detailed metrics are provided in the supplementary materials.

### 4.4 Batch and Multi-omic integration

In the multi-batch integration task, we first matched the genes in the downstream task data with the SC-MAMBA2 vocabulary to select the shared genes, and then identified the highly variable genes across multiple batches. Similar to the cell type annotation task, we did not filter out genes with zero expression in each cell. We then used the [CLS] token as the representation for each cell, added batch tokens, and applied an adversarial domain adaptation approach[48] to map these tokens into the same space, fine-tuning SC-MAMBA2. The fine-tuned [CLS] token was obtained as the embedding for each cell.

For single-cell multi-omic integration tasks, we retained the pretrained single-cell transcriptomics latent weights and reinitialized weights for other modalities (e.g., ATAC-seq, protein). Using a preprocessing approach similar to that for multi-batch integration, each cell obtained a [CLS] token as its representation, with an added modality token to indicate the data type for each cell. We applied an adversarial domain adaptation approach to map these representations into the same space, fine-tuning SC-MAMBA2 for multi-omic data integration.

We used scib[33] to evaluate the integration performance, where AvgBio represents the average of ARI, NMI, and ASW cell scores. All detailed metrics are provided in the supplementary materials.

### 4.5 Perturbation and reverse perturbation

For the perturbation prediction task, SC-MAMBA2 utilizes all genes from the downstream dataset without filtering out zero-expression genes and uses log-normalized expression values as input instead of binned values to enhance the accuracy of predicting absolute expression values post-perturbation. Additionally, a binary condition token is introduced at each input gene position to indicate whether the gene has been perturbed. SC-MAMBA2 initializes the model using parameters from the embedding layers and Biamba layers of the pretrained model and fine-tunes it in a supervised manner by minimizing the difference between predicted and actual gene expression, subsequently applying it to the test set.

The evaluation of the perturbation prediction task includes four settings. For single-gene perturbation, the model predicts single-gene perturbations not seen in the training set (Single Unseen). In the dual-gene perturbation tasks, both genes are unseen in the training set (Seen 0), one gene appears in the training set (Seen 1), or both genes appear in the training set (Seen 2). Model evaluation calculates the correlation between predicted and actual expression. All detailed metrics are provided in the supplementary materials.

## Supporting information

SUPPLEMENTARY MATERIAL OF SC-MAMBA2

## Data availability

The pretraining datasets were sourced from the CELLxGENE census (release version 1 July 2024, accessible at https://chanzuckerberg.github.io/cellxgene-census/python-api.html and https://cellxgene.cziscience.com/). For annotation tasks, the MS dataset was retrieved from the Gene Expression Atlas (https://www.ebi.ac.uk/gxa/sc/experiments/E-HCAD-35), while the myeloid dataset is publicly available via GEO (GSE154763). The processed human pancreas dataset was obtained from https://github.com/JackieHanLab/TOSICA. Reference mapping utilized the Lung-Kim dataset from the Curated Cancer Cell Atlas (https://www.weizmann.ac.il/sites/3CA/lung), and the processed COVID-19 dataset was accessed from https://github.com/theislab/scarches-reproducibility. Perturbation prediction tasks used the Norman and Adamson datasets from Harvard Dataverse (https://dataverse.harvard.edu/api/access/datafile/6154020 and https://dataverse.harvard.edu/api/access/datafile/6154417). The Replogle dataset was retrieved from https://gwps.wi.mit.edu/. For batch integration, the PBMC 10k dataset was accessed via scVI tools (https://scvi-tools.org/), while the perirhinal cortex dataset was sourced from the CELLxGENE Human Brain Cell Atlas v1.0 (https://cellxgene.cziscience.com/collections/283d65eb-dd53-496d-adb7-7570c7caa443). The multi-omics integration task utilized the 10x Multiome PBMC dataset (https://scglue.readthedocs.io/en/latest/data.html), the BMMC dataset (GSE194122) from GEO.

## Code availability

The codebase for SC-MAMBA2 is publicly available at https://github.com/xtalpi-xic/SC-MAMBA2

## Acknowledgments

We are deeply grateful to the XtalPI Innovation Center (XIC) for the financial support and provision of essential computational resources that made this research possible. Special thanks go to Dr. Zhenghui Wang from XIC for her invaluable insights and constructive feedback throughout the study. We also appreciate the collaborative support of our partner, Gnosis Neurodynamics, whose inspiring discussions significantly contributed to the development of this work. Lastly, we acknowledge the Chan Zuckerberg Initiative for providing the CELLxGENE dataset, as well as all other publicly available datasets utilized in this study, which have been instrumental in advancing our research and the field of single-cell transcriptomics.

