## SUPPLEMENTARY MATERIAL OF SC-MAMBA2 for "SC-MAMBA2: Leveraging State-Space Models for Efficient Single-Cell Ultra-Long Transcriptome Modeling"

### ABSTRACT

The rapid advancement of single-cell sequencing technology has significantly deepened our understanding of cellular heterogeneity, yet it concurrently presents substantial challenges for the unified modeling of single-cell data. Simultaneously, pre-trained foundation models have achieved notable success in domains such as natural language processing and image analysis. However, extending these models to accommodate ultra-long single-cell transcriptome sequences, characterized by an extensive number of genes, remains a formidable task. In this study, we introduce SC-MAMBA2, based on the MAMBA2 architecture, meticulously designed with a bidirectional modeling approach tailored for single-cell transcriptomic data. As the first single-cell foundation model to integrate state-space models (SSMs) underlying MAMBA2 architecture, SC-MAMBA2 features over 625 million parameters, covers more than 60,000 genes, and was pre-trained on a dataset of over 57 million cells, making it the most comprehensive solution for processing ultra-long transcriptome sequences. Extensive benchmarking across a diverse array of downstream tasks consistently demonstrates that SC-MAMBA2 surpasses state-of-the-art models, delivering superior accuracy and enhanced computational efficiency.

***Keywords*** single-cell transcriptomics · state-space model · SC-MAMBA2 · ultra-long sequences · gene expression analysis · large-scale modeling · computational biology

[*]These authors contributed equally to this work.
[†]This work was completed during an internship at XtalPi.

Table 1: Cell type annotation results on the hPancreas dataset

| Model | Accuracy | Precision | Recall | MacroF1 |
|---|---|---|---|---|
| TOSICA | 0.96 | 0.661 | 0.681 | 0.656 |
| scGPT | 0.968 | 0.735 | 0.725 | 0.718 |
| scMamba | 0.975 | 0.776 | 0.768 | 0.761 |

Table 2: Cell type annotation results on the Myeloid dataset

| Model | Accuracy | Precision | Recall | MacroF1 |
|---|---|---|---|---|
| TOSICA | 0.488 | 0.316 | 0.276 | 0.275 |
| scGPT | 0.642 | 0.366 | 0.347 | 0.346 |
| scMamba | 0.644 | 0.381 | 0.366 | 0.364 |

Table 3: Cell type annotation results on the Multiple Sclerosis dataset

| Model | Accuracy | Precision | Recall | MacroF1 |
|---|---|---|---|---|
| TOSICA | 0.758 | 0.664 | 0.585 | 0.578 |
| scGPT | 0.856 | 0.729 | 0.72 | 0.703 |
| scMamba | 0.869 | 0.732 | 0.73 | 0.723 |

Table 4: Integration results on the PBMC 10K dataset

| **Model** | **AvgBIO** | **NMIcell** | **ARIcell** | **ASWcell** |
|---|---|---|---|---|
| scMamba | 0.8360 | 0.8508 | 0.8978 | 0.7594 |
| scGPT | 0.8210 | 0.8500 | 0.8730 | 0.7400 |
| scVI | 0.7530 | 0.8190 | 0.8470 | 0.5920 |
| Seurat v4 | 0.7240 | 0.8080 | 0.7220 | 0.6410 |
| Harmony | 0.7840 | 0.8600 | 0.9020 | 0.5910 |

Table 5: Integration results on the 10X Multiome PBMC dataset

| **Model** | **AvgBIO** | **NMIcell** | **ARIcell** | **ASWcell** |
|---|---|---|---|---|
| scMamba | 0.762 | 0.826 | 0.827 | 0.634 |
| scGPT (fine-tuned) | 0.758 | 0.807 | 0.822 | 0.645 |
| scGLUE | 0.747 | 0.815 | 0.806 | 0.619 |
| Seurat v4 | 0.722 | 0.784 | 0.691 | 0.691 |

Table 6: Integration results on the BMMC dataset

| **Model** | **AvgBIO** | **NMIcell** | **ARIcell** | **ASWcell** |
|---|---|---|---|---|
| scMamba | 0.701 | 0.787 | 0.733 | 0.582 |
| scGPT | 0.697 | 0.783 | 0.725 | 0.582 |
| Seurat v4 | 0.642 | 0.707 | 0.47 | 0.594 |

Table 7: Perturbation results for Single Unseen

| Metric | Gears | scFoundation | scGPT | scMamba |
| --- | --- | --- | --- | --- |
| pearson_delta | 0.3605 | 0.4226 | 0.4167 | 0.4073 |
| pearson_delta_de | 0.5640 | 0.4229 | 0.5566 | 0.4924 |

Table 8: Perturbation results for Seen 0

| Metric | Gears | scFoundation | scGPT | scMamba |
| --- | --- | --- | --- | --- |
| pearson_delta | 0.5381 | 0.5381 | 0.5706 | 0.5906 |
| pearson_delta_de | 0.7749 | 0.4280 | 0.6721 | 0.8425 |

Table 9: Perturbation results for Seen 1

| Metric | Gears | scFoundation | scGPT | scMamba |
| --- | --- | --- | --- | --- |
| pearson_delta | 0.5634 | 0.4452 | 0.6385 | 0.6534 |
| pearson_delta_de | 0.8493 | 0.4811 | 0.7705 | 0.7550 |

Table 10: Perturbation results for Seen 2

| Metric | Gears | scFoundation | scGPT | scMamba |
|---|---|---|---|---|
| pearson_delta | 0.5774 | 0.5399 | 0.7447 | 0.8004 |
| pearson_delta_de | 0.6758 | 0.7074 | 0.8896 | 0.9065 |

Table 11: Perturbation results (Average)

| Metric | Gears | scFoundation | scGPT | scMamba |
|---|---|---|---|---|
| pearson_delta | 0.5098 | 0.4865 | 0.5926 | 0.6129 |
| pearson_delta_de | 0.7160 | 0.5099 | 0.7222 | 0.7491 |
